# *SLC35A2* modulates paramyxovirus fusion events during infection

**DOI:** 10.1101/2024.08.27.609835

**Authors:** Yanling Yang, Yuchen Wang, Danielle E. Campbell, Heng-Wei Lee, Leran Wang, Megan Baldridge, Carolina B. López

## Abstract

Paramyxoviruses are significant human and animal pathogens that include mumps virus (MuV), Newcastle disease virus (NDV) and the murine parainfluenza virus Sendai (SeV). Despite their importance, few host factors implicated in paramyxovirus infection are known. Using a recombinant SeV expressing destabilized GFP (rSeVC^dseGFP^) in a loss-of-function CRISPR screen, we identified the CMP-sialic acid transporter (CST) gene *SLC35A1* and the UDP-galactose transporter (UGT) gene *SLC35A2* as essential for paramyxovirus infection. *SLC35A1* knockout (KO) cells showed significantly reduced binding and infection of SeV, NDV and MuV due to the lack of cell surface sialic acids, which act as their receptors. However, *SLC35A2* KO cells revealed unknown critical roles for this factor in virus-cell and cell-to-cell fusion events during infection with different paramyxoviruses. While the UGT was essential for virus-cell fusion during SeV entry to the cell, it was not required for NDV or MuV entry. Importantly, the UGT promoted the formation of larger syncytia during MuV infection, suggesting a role in cell-to-cell virus spread. Our findings demonstrate that paramyxoviruses can bind to or enter A549 cells in the absence of canonical galactose-bound sialic-acid decorations and show that the UGT facilitates paramyxovirus fusion processes involved in entry and spread.

## Introduction

The Paramyxovirus family includes the major human and animal pathogens measles virus (MV), mumps virus (MuV), human parainfluenza virus (hPIV), Newcastle disease virus (NDV) and the highly pathogenic zoonotic Hendra (HeV) and Nipah (NiV) viruses imposing a significant burden on global public health, while also causing substantial economic losses [1, 2].

Paramyxoviruses are single-stranded negative-sense RNA enveloped viruses containing a fusion (F) and an attachment glycoprotein on their surface. These glycoproteins are essential for virus entry and infection. Attachment proteins vary across the different genera of paramyxoviruses, and can be either the glycoprotein (G), the hemagglutinin (H), or the hemagglutinin-neuraminidase (HN). The *Avulavirus* (e.g., NDV), *Rubulavirus* (e.g., MuV), and *Respirovirus* (e.g., SeV) genera attach to the cell surface through the virus HN protein that binds sialic acid-containing cell surface molecules [3]. Sialic acids also serve as attachment receptors for many other viruses, including influenza virus, reovirus, adenovirus, and rotavirus [4]. After the virus attaches to the host cell, the F protein undergoes a conformational change that triggers the fusion of the host cell and viral membranes. Virus-cell membrane fusion leads to the release of the viral ribonucleoprotein complex into the cytosol, allowing for viral replication and transcription to occur [5]. In some cases, the virus enters and fuses with the endosomal membrane [6, 7]. In addition, the F protein can facilitate cell-to-cell fusion and syncytia formation, for example during MuV infection [8–10].

Sialic acids are bound to carbohydrate chains on glycoproteins and glycolipids in the Golgi apparatus via different glycosidic linkages. The most common linkage types are α2,3-linkage to a galactose residue, α2,6-linkage to a galactose residue, α2,6-linkage to an N-acetylgalactosamine residue, and α2,8-linkage to another sialic acid moiety on a glycan [4]. Sialic acids and galactose are transported into the Golgi by the CMP-sialic acid transporter (CST) and the UDP-galactose transporter (UGT) encoded by SLC35A1 and *SLC35A2*, respectively. CST facilitates the assembly of sialic acid onto glycoproteins and glycolipids [11]. SeV and MuV are reported to only use α2,3-linked sialic acid to attach to cells [12–15], while NDV can bind to both α2,3-linked and α2,6-linked sialic acids [16]. All reported paramyxovirus receptors involve sialic acids linked to a galactose, suggesting that this glycan motif may be essential for sialic acid-dependent virus infection.

Targeting host factors essential to the viral lifecycle is one promising avenue for antiviral drug development [17, 18]. Unfortunately, the list of known cellular host factors and their importance in modulating the paramyxovirus lifecycle is relatively sparse when compared to other viruses such as influenza virus and coronavirus [19–22]. Even less is known about the common and divergent host protein requirements among different paramyxoviruses. Given the significance of paramyxoviruses in disease and the lack of clear candidates for a host-directed antiviral drug design, unbiased and high-throughput screening for host factor dependencies remains a necessary research objective for this virus family.

In this work, we used the murine paramyxovirus Sendai virus (SeV) which causes respiratory infection in mice and is widely used as a model paramyxovirus [23–28], to perform CRISPR-Cas9-based screenings for essential pro-viral host factors. We leveraged a novel recombinant SeV strain expressing a destabilized eGFP (dseGFP) reporter that allowed for sensitive measurements of viral genome replication and transcription within the infected cell, permitting more accurate analysis of the CRISPR-Cas9 knockout (KO) library screening results. Consistent with several published screens for other sialic-acid dependent RNA viruses, we found that the top essential pro-viral genes included *SLC35A1* [29–32] and *SLC35A2* [32, 33]. *SLC35A1* serves as an essential gene for the expression of the virus attachment receptors. In contrast, we discovered that UGT, in addition to contributing to virus attachment, plays independent roles in paramyxovirus virus-cell and cell-cell fusion processes.

## Results

### CRISPR knock-out screen identifies *SLC35A1* and *SLC35A2* as essential factors for Sendai virus infection

To identify host factors essential for paramyxovirus infection, we developed genome-wide CRISPR KO libraries in A549 cells to screen for infection with the model virus SeV (Fig 1A). Cas9-stable A549 cells were generated by transduction with a lentivirus expressing Cas9. Several single cell clones of A549-Cas9 cells were selected based on the expression of Cas9 as determined by western blot (Fig S1A). The Cas9 activity of the clones was then confirmed by an eGFP knockout assay where higher Cas9 activity results in a lower percentage of GFP positive cells (Fig S1B) [34]. The A549-Cas9 single cell clone 1 with the highest Cas9 efficiency (Fig S1C) was selected and transduced at a low multiplicity of infection (MOI) of 0.3, with the Human CRISPR KO lentiviral single guide (sg) RNA Library Brunello [35] followed by puromycin selection.

**Fig 1.**
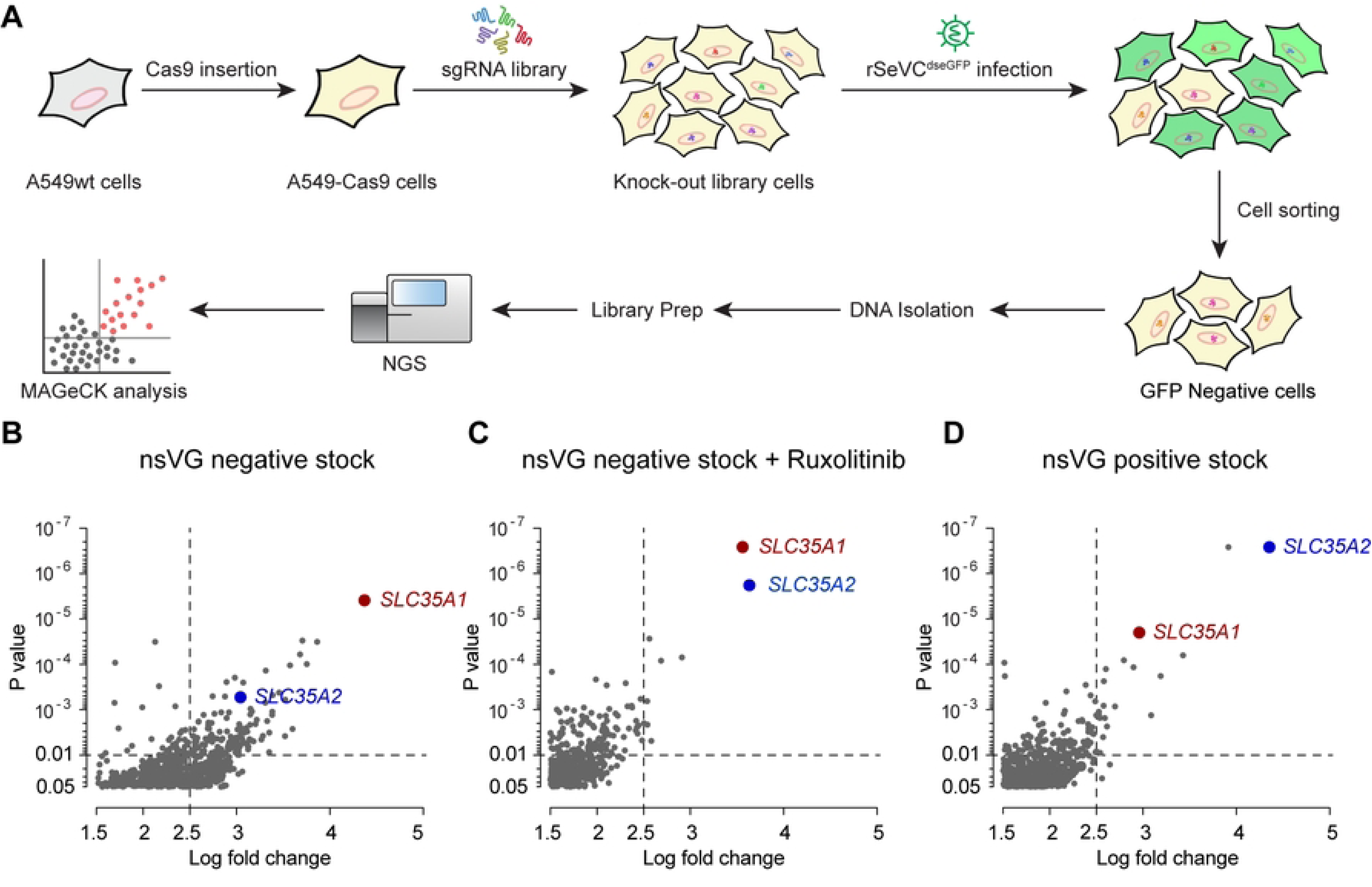
CRISPR screen workflow and sgRNA enrichment analysis. (A) Summary of the CRISPR screen workflow, from the generation of the A549-Cas9 stable cell line to the sequencing and MAGeCK analysis. (B-D) Scatter plots showing the enrichment of sgRNAs in GFP-negative cells at 24 hpi relative to control unsorted mock cells (P value < 0.05, log2 fold change > 1.5). Cells were infected with rSeV^dseGFP^ nsVG negative stock at an MOI of 10 or rSeV^dseGFP^ nsVG positive stock at an MOI of 3. Differences in enrichment were calculated as log2-normalized fold change. *SLC35A1* and *SLC35A2* sgRNAs were significantly enriched in all three independent screenings: using nsVG negative virus stock (B), nsVG negative virus stock with 5uM Ruxolitinib treatment (C), and nsVG positive virus stock (D).

For screening, we generated a recombinant SeV expressing a destabilized eGFP (rSeVC^dseGFP^). This virus was generated by inserting a destabilized eGFP (dseGFP) between SeV NP and P genes (Fig S2A). The destabilization is due to a fused proline-glutamate-serine-threonine-rich (PEST) peptide to eGFP, which reduces the half-life of GFP from 20 hours to 2 hours and cause a 90% signal loss [36, 37]. As shown in Fig S2B, the rSeVC^dseGFP^ did not show signs of attenuation in virus titer (10^8.28^ vs 10^8.35^ TCID_50_/ml) but exhibited lower eGFP intensity compared with rSeVC^eGFP^ in infected A549 cells. To identify host factors regulating infection regardless of the antiviral response, we performed three screens using different immunostimulatory conditions. First, transduced cells were infected with either rSeVC^dseGFP^ nonstandard viral genomes (nsVG)-negative stocks in the absence or presence of the JAK/STAT signaling inhibitor Ruxolitinib. Stocks without nsVGs lack strong immunostimulatory molecules [38], whereas drug treatment precludes interferon signaling. Second, another batch of transduced cells were infected with rSeVC^dseGFP^ stock with a high content of immunostimulatory nsVGs (nsVG positive), which induce strong immune responses [39]. Among the subset of sgRNAs that were enriched in the GFP-negative cell population in all three independent screenings relative to the control (S1-S3 Tables), we identified the genes *SLC35A1* and *SLC35A2* encoding the CMP-sialic acid transporter and the UDP-galactose transporter as significantly enriched (log fold change >2.5, p<0.01) (Fig 1C).

### *SLC35A1* and *SLC35A2* are essential for SeV infection

The *SLC35A1* gene encodes the CMP-sialic acid transporter (CST) necessary for the sialylation of proteins and lipids [11]. The *SLC35A2* gene encodes the UDP-galactose transporter (UGT) which is required not only for the galactosylation of N- and O-glycans on glycoproteins but also for the synthesis of galactosylceramide and galactosyl diglyceride [11]. CST and UGT are found in the membrane of the Golgi apparatus and transport CMP-sialic acid and UDP-galactose from the cytosol into Golgi vesicles for the generation of glycans (Fig 2A, adapted from [11]). The terminal sugar chains of sialylated glycoproteins and gangliosides, such as GD1a and GQ1b, that act as SeV receptors are shown in Fig 2B. To validate the functional significance of *SLC35A1* and *SLC35A2* during SeV infection, these genes were disrupted in A549-Cas9 cells using sgRNAs. A control cell line was made by transducing a scramble sgRNA that did not target any specific host gene. Transduced cells were then selected with puromycin followed by singe cell cloning. We then tested for the presence of surface sialic acids and galactose in the KO cell lines by staining with the lectins Sambucus Nigra Agglutinin (SNA) and Erythrina Cristagalli Lectin (ECL) to detect cell-surface sialic acid and galactose, respectively as previously described [40, 41] (Fig 2C). As shown in Fig 2D, *SLC35A1* KO cells lack cell surface sialic acid while having more exposed galactose [41]. *SLC35A2* KO cells lack galactose in the cell surface and as expected since most of terminal sialic acid are linked to galactoses, have a significantly reduced level of sialic acid.

**Fig 2.**
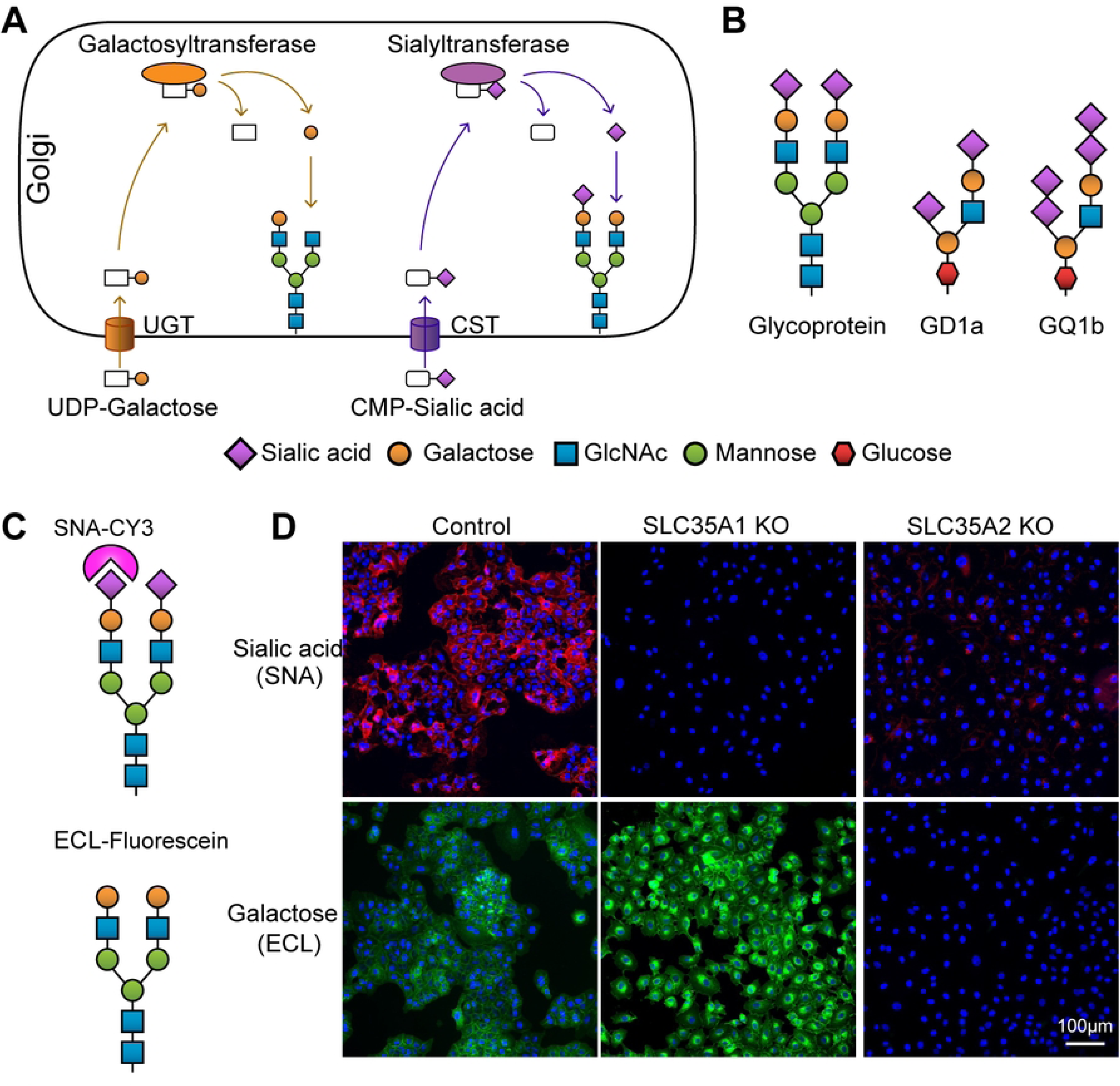
Roles of *SLC35A1* and *SLC35A2* and lectin staining analysis in A549 KO cells. (A) Schematic of glycosylation pathways in the Golgi apparatus, showing the roles of the *SLC35A1* gene encoding CMP-sialic acid transporter (CST) and the *SLC35A2* gene encoding UDP-galactose transporter (UGT). Galactosyltransferase and sialyltransferase enzymes add galactose and sialic acid residues to glycans, respectively. (B) Examples of SeV receptors: glycoprotein and gangliosides GD1a and GQ1b, illustrating the incorporation of sialic acid and galactose residues. (C) Diagram of lectin staining: SNA (Sambucus nigra agglutinin) binds to cell surface sialic acid, while ECL (Erythrina cristagalli lectin) binds to galactose. (D) Analysis of sialic acid and galactose expression by lectin staining. Control, *SLC35A1* KO, and *SLC35A2* KO cells were fixed and stained with lectins SNA or ECL specific for sialic acid or galactose and analyzed by fluorescence microscopy. The images show the distribution of sialic acid (magenta) and galactose (green) residues. Scale bar lengths are indicated.

We then used a SeV reporter virus expressing eGFP (rSeVC^eGFP^) to directly assess the impact of *SLC35A1* or *SLC35A2* during infection. As a control, vesicular stomatitis virus (VSV) was used as it does not depend on sialic acid for entry. We looked for GFP expression at 24 hpi with an MOI of 1.5 for SeV and an MOI of 0.015 for VSV as a readout of infection and virus replication. Absence of *SLC35A1* and *SLC35A2* resulted in loss of infectivity in most cells, with only a few cells showing viral replication (Fig. 3A). We confirmed absence of SeV replication in *SLC35A1* KO cells and drastically reduced replication in *SLC35A2* KO cells by evaluating SeV NP mRNA expression by qPCR (Fig. 3B). In contrast, VSV infection proceeded normally in the absence of *SLC35A1* or *SLC35A2*. To exclude the possibility of other defects that may result in the restriction of SeV infection in the KO cells, we complemented *SLC35A1* KOs with DNA expressing *SLC35A1*-GFP and *SLC35A2* KOs with cDNA expressing *SLC35A2*-GFP. At 24 hpi with 3 MOI of rSeVC^miRF670^, we observed recovered SeV replication in complemented KOs (Fig 3C and 3D). Infection using high MOIs of rSeVC^dseGFP^ or rSeVC^eGFP^ showed more cells infected in *SLC35A1* KO cells, and an even larger number in *SLC35A2* KO cells at 24hpi, but in both cases the percentage of cells infected was significantly less than controls, suggesting that SeV can enter more cells when used at high MOIs independent of *SLC35A1* and *SLC35A2* but with limited spread (Fig S3). Overall, these data confirmed the critical, yet not completely overlapping, roles of *SLC35A1* and *SLC35A2* during SeV infection.

**Fig 3.**
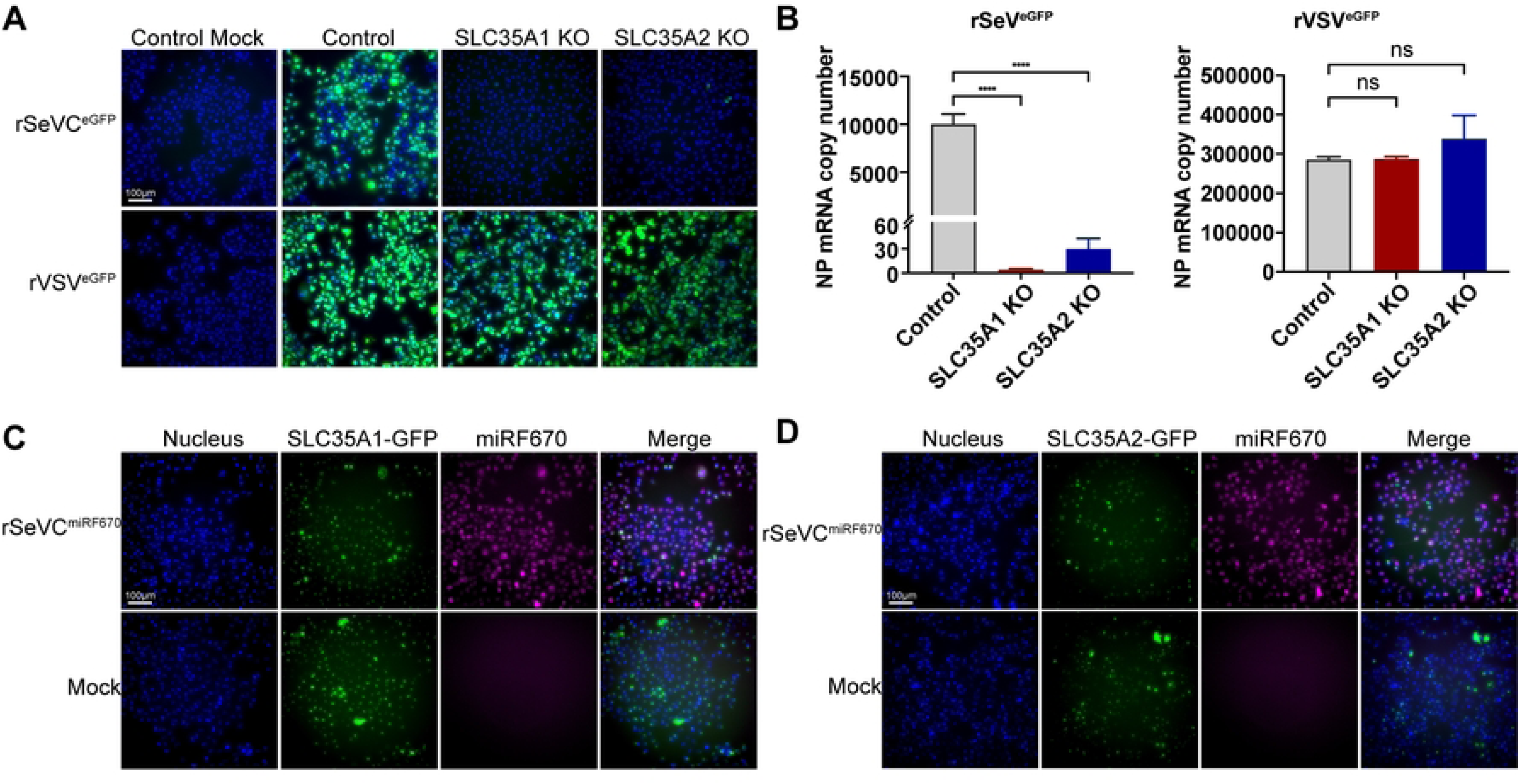
*SLC35A1* and *SLC35A2* are essential for optimal SeV infection. (A) Fluorescence images showing GFP expression in control, *SLC35A1* KO, and *SLC35A2* KO A549 cells infected with rSeVC^eGFP^ at an MOI of 1.5 or rVSV^eGFP^ at an MOI of 0.015, 24hpi. Scale bar lengths are indicated. (B) Quantification of NP mRNA copy number in KO cells infected with rSeVC^eGFP^ or rVSV^eGFP^. Control cells, *SLC35A1* KOs, and *SLC35A2* KOs were infected with rSeVC^eGFP^ at an MOI of 1.5 or rVSV^eGFP^ at an MOI of 0.015 and cells were collected at 24hpi followed by relative qPCR analysis. Expression of mRNA calculated relative to the housekeeping index with *GAPDH* and *β-actin*. Data represent the mean of three independent experiments. ****: p<0.0001, ns: not significant. (C, D) Fluorescence images showing complementation of KO cells with either (C) *SLC35A1*-GFP or (D) *SLC35A2*-GFP. Complemented cells were infected with rSeVC^miRF670^ at a MOI of 3, and images were analyzed at 24hpi. The nucleus was stained with Hoechst (Blue). Scale bar lengths are indicated.

### *SLC35A1* and *SLC35A2* differentially impact infection with several paramyxoviruses

To assess whether the observed non-overlapping functions of *SLC35A1* and *SLC35A2* were maintained during infection with other paramyxoviruses, we infected single and double KO A549 cells with the *Respirovirus* rSeVC^eGFP^, the *Avulavirus* rNDV^eGFP^, or the *Rubulavirus* MuV at an MOI of 1.5 and look for either GFP expression (SeV and NDV) or stained for MuV NP at 24 hpi. The double KO cell was made by transducing *SLC35A1* KO cells with *SLC35A2* sgRNA and it was confirmed by lectin staining (Fig 4A). In all cases, the absence of *SLC35A1* or both *SLC35A1* and *SLC35A2* resulted in loss of infectivity in most cells, with only a few cells showing viral replication. However, for NDV and MuV infections, the absence of *SLC35A2* alone resulted in an intermediate number of infected cells showing more infected cells than the double KO cells but significantly less than control cells (Fig. 4B and 4C). These data demonstrate that *SLC35A2* has non-redundant functions with *SLC35A1* during paramyxovirus entry and spread and suggest a differential impact of *SLC35A2* during infection with different paramyxoviruses.

**Fig 4.**
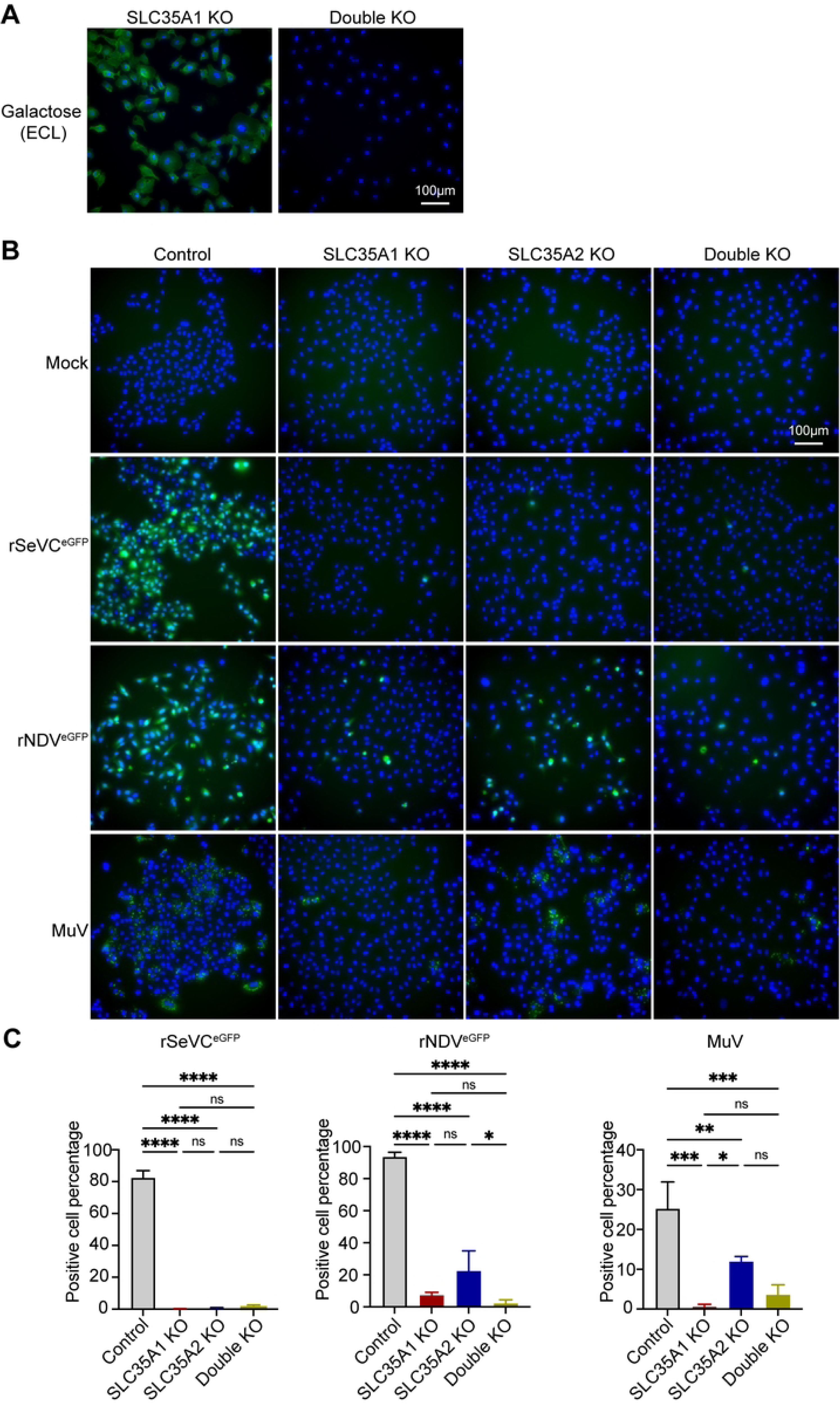
Impact of *SLC35A1* and *SLC35A2* single KOs or double KOs in infection with different paramyxoviruses. (A) Analysis of galactose expression in *SLC35A1*/*SLC35A2* double KO cells by lectin staining. *SLC35A1* KO cells, and double KO cells were fixed and stained with ECL followed by fluorescence microscopy. The images show the distribution of galactose (green) residues. The nucleus was stained with Hoechst 33342 (Blue). Scale bar lengths are indicated. (B) Control and KO cell lines were infected with rSeVC^eGFP^, rNDV^eGFP^, or MuV at an MOI of 1.5. Fluorescence images showing GFP expression (rSeVC^eGFP^ and rNDV^eGFP^) or NP staining (MuV) in control, *SLC35A1* KO, *SLC35A2* KO, and double KO cells at 24hpi. Scale bar lengths are indicated. (C) Quantification of infected cells in (B). Significance was calculated with an ordinary one-way ANOVA. ns: not significant, *: p<0.05, **: p<0.01, ***: p<0.001, ****: p<0.0001.

### *SLC35A2* differentially impacts SeV, NDV, and MuV infection and spread in A549 cells

To further investigate the impact of *SCL35A2* on paramyxovirus infection and spread, we followed the infection in *SCL35A2* KO cells through a 4-day infection period (Fig. 5). Interestingly, infections in *SLC35A2* KO cells displayed varied phenotypes across these viruses. As shown before, SeV infection was drastically reduced to one or two cells per image (5X magnification) and the virus did not spread throughout the time course (Fig. 5A-B). NDV could infect a larger proportion of *SLC35A2* KO cells before the infected cells died (Fig S4), but again, there was no evidence of virus spread, compared to NDV replication in control cells (Fig. 5C-D). In contrast, MuV infected and spread well in *SLC35A2* KO cells as evidenced by staining for the virus NP (Figure 5E-F). Taken together, these data indicate that *SLC35A2* plays differential roles in the infection and spread of different paramyxoviruses during infection of A549 cells.

**Fig 5.**
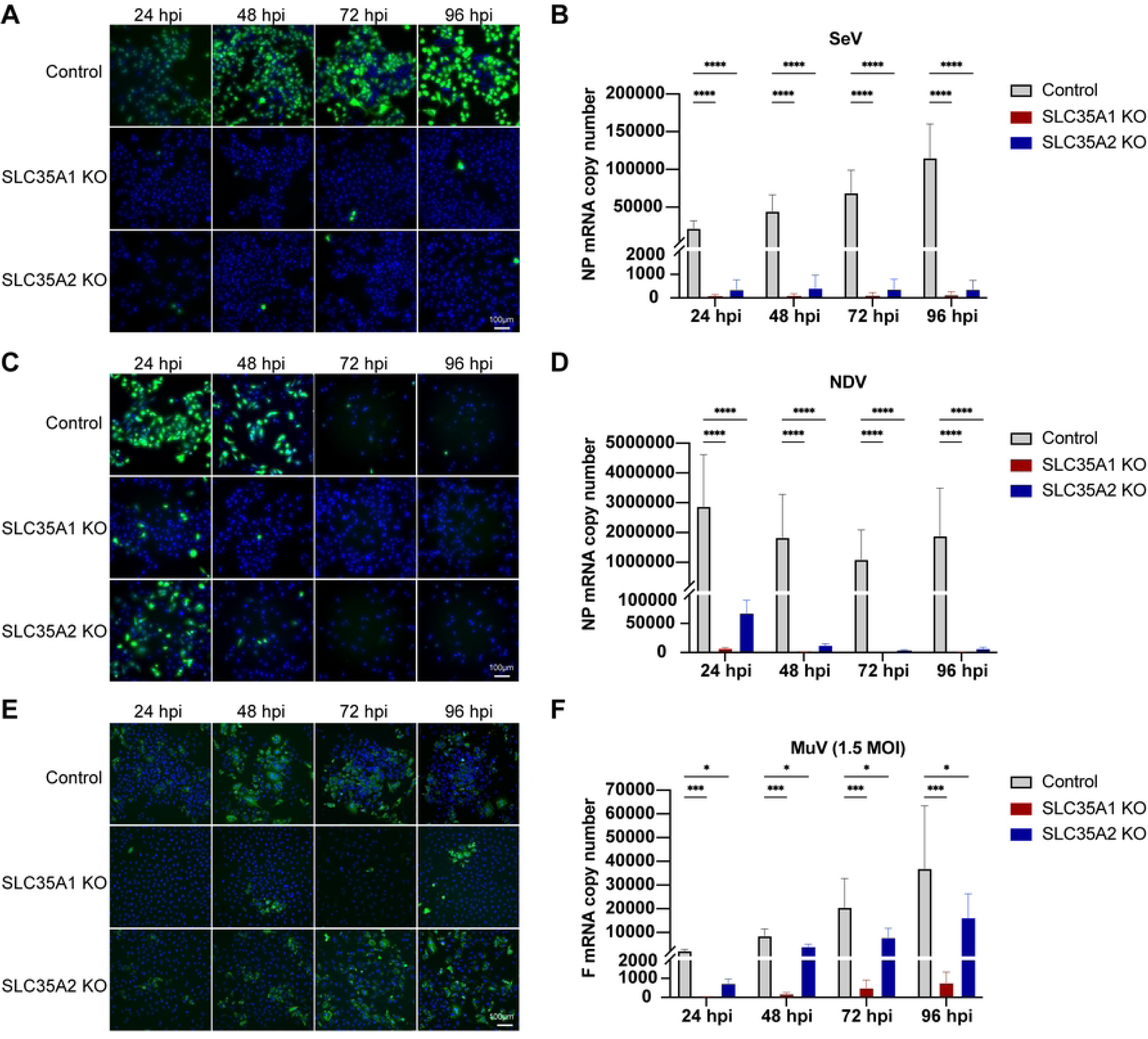
Time course infection with paramyxoviruses in *SLC35A1* and *SLC35A2* KO A549 cells. (A, C and E) Fluorescence (A and C) or immunofluorescence (E) images of control, *SLC35A1* KO, and *SLC35A2* KO cells infected with rSeVC^eGFP^ (A), rNDV^eGFP^ (C), or MuV (E) at an MOI of 1.5. Images were analyzed at 24, 48, 72, and 96hpi. The nucleus was stained with Hoechst 33342 (Blue), green fluorescence indicates viral infection represented by eGFP or MuV NP staining. Data shown represent one of three independent experiments. (B, D, and F) Under the same infection conditions as A, C, and E, cellular RNA was collected at 24-96hpi and analyzed by relative qPCR. Expression of mRNA was calculated relative to the housekeeping index. Data represent the mean of three independent experiments. Significance was calculated with a two-way ANOVA. ns: not significant, *: p<0.05, ***: p<0.001, ****: p<0.0001

### *SLC35A2* is essential for virus-cell fusion during SeV infection

We next focused on investigating where *SLC35A2* played a critical role during the virus infection cycle. Based on the organization of the terminal sugar chains of sialylated glycoproteins and the apparent requirement for galactose for sialic acid proximal binding (Fig. 2B), we began by evaluating whether *SLC35A2* impacts the attachment of SeV to the cell surface. As expected, *SLC35A1* KO cells exhibited robust restriction of SeV binding evidenced by consistently negative cell surface HN staining in both infected and non-infected groups after co-incubation of virus and cells for 1 hours at 4°C. However, incubation of SeV with *SLC35A2* KO cells under the same conditions resulted in positive cell surface HN staining, albeit lower than control cells (Fig. 6A), suggesting that the impact of *SLC35A2* on paramyxovirus infection extends beyond the virus binding step.

**Figure 6.**
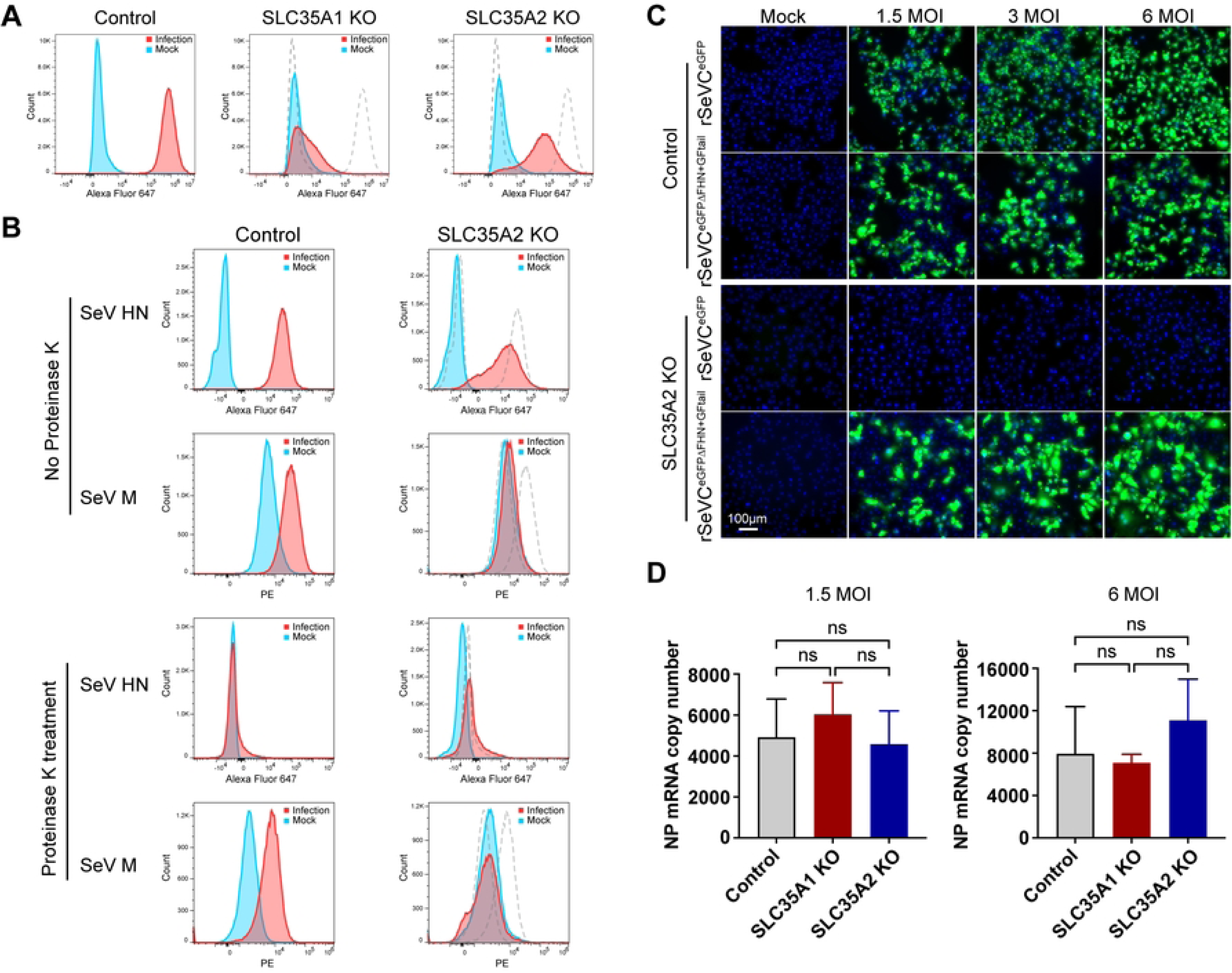
S*L*C35A2 is essential for SeV-cell fusion. (A) A549 control, *SLC35A1* KOs, or *SLC35A2* KOs incubated with SeV at an MOI of 30 (infection), or with infection media (mock), at 4°C for 1 hour, followed by staining using an anti-HN antibody (Alexa Fluor 647). The dashed lines in the middle and right panels represent the histograms of the control cell line. Data shown represent one of three independent experiments. (B) A549 control cells or *SLC35A2* KOs were incubated with rSeV-M-HA at an MOI of 400 (infection), or with infection media (Mock), at 37°C for 3 hours, followed by 70 minutes of proteinase K treatment (0.5mg/ml) at 37℃. The cells were then fixed and permeabilized. Anti-HA antibody was used to detect intracellular M protein, while HN protein was detected as an extracellular control. The dashed lines in the right panels represent the histograms of the control cell line. Data shown represent one of three independent experiments. (C) rSeVC^eGFP^ and rSeVC^eGFPΔFHN+GFtail^ were used to infect A549 control cells, or *SLC35A2* KOs at MOIs of 1.5, 3, or 6. Images were analyzed at 24hpi. The nucleus was stained with Hoechst 33342 (blue). Green fluorescence indicates reporter gene expression. Scale bar lengths are indicated. (D) rSeVC^eGFPΔFHN+GFtail^ was used to infect A549 control cells, or *SLC35A2* KOs at MOIs of 1.5 or 6. Cellular RNA was collected 24 hours later and analyzed by qPCR. Expression of mRNA was calculated relative to the housekeeping index. Data represent the mean of three independent experiments. Significance was calculated with a two-way ANOVA. ns: not significant.

To directly test the impact of *SLC35A2* in the SeV virus-cell fusion process, we tested for intracellular detection of the SeV internal M (matrix) protein as a marker for fusion [42] using a recombinant SeV rSeV-M-HA [43] in which the M protein is fused with an HA tag. In brief, SeV was incubated with cells at 37°C for 3 hours to allow virus entry and fusion. The cells were then treated with Proteinase K followed by fixation, permeabilization, and staining of SeV M and envelope HN proteins. Proteinase K treatment was used to remove cell surface attached viral particles and the HN protein staining was used as a control for the presence of virus attached to the outside of the cells. As shown in Figure 6B, SeV HN was detected in both cell lines, similar to Figure 6A, but not after proteinase K treatment. However, the M protein was detected in the control cells but not in *SLC35A2* cells, regardless of proteinase K treatment, suggesting that *SLC35A2* is essential for efficient fusion of the virus and cell membrane during virus entry.

Lastly, to confirm that *SLC35A2* did not directly affects SeV genome replication and transcription, we took advantage of the recombinant reporter SeV virus rSeVC^eGFPΔFHN+GFtail^. This virus was made by removing the original SeV F and HN and inserting a chimeric VSV glycoprotein G fused with the C-terminal tail of SeV F (GFtail) thus replicating as SeV but entering the cells as VSV (Fig. S5A). As expected, rSeVC^eGFPΔFHN+GFtail^ can infect *SLC35A1* KO cells using the VSV GFtail protein for entry (Fig S5B). Then we asked whether rSeVC^eGFPΔFHN+GFtail^ can replicate without *SLC35A2*. Although both viruses infect control cells to a similar degree, rSeVC^eGFP^ was unable to infect and replicate in *SLC35A2* KO cells but rSeVC^eGFPΔFHN+GFtail^ infected and spread normally in these cells (Fig. 6C). In addition, SeV transcription measured by SeV NP mRNA expression showed that *SLC35A1* and *SLC35A2* deletions have no effect on the viral polymerase activity (Fig. 6D). These data demonstrate that *SLC35A2* is not essential for virus binding or viral genome replication and transcription but is critical for virus-cell fusion during SeV infection.

### *SLC35A2* is implicated in syncytia formation during MuV infection

As MuV infected and transcribed in *SLC35A2* KO cells (Fig. 4E-F), we next asked whether *SLC35A2* impacted other steps of the MuV replication cycle. As shown in Fig. 7A and 7B, there was no significant difference in the virus production of infectious viral particles between control and *SLC35A2* KO cells infected at MOI of 1.5 or 15, indicating that *SLC35A2* does not impact MuV infectious particle production. Interestingly, we noticed less syncytia formation in *SLC35A2* KO cells at both low and high MOI. To confirm this difference, we quantified the number of syncytia formed under low and high MOI conditions at 48 hpi. We defined an NP-positive large cell with more than three nuclei as syncytia. As shown in Fig. 7C-E, *SLC35A2* KO cells formed less syncytia compared to control cells at both MOIs, suggesting that *SLC35A2* is implicated in MuV-induced syncytia formation suggesting a role in cell-to-cell virus spread.

**Fig 7.**
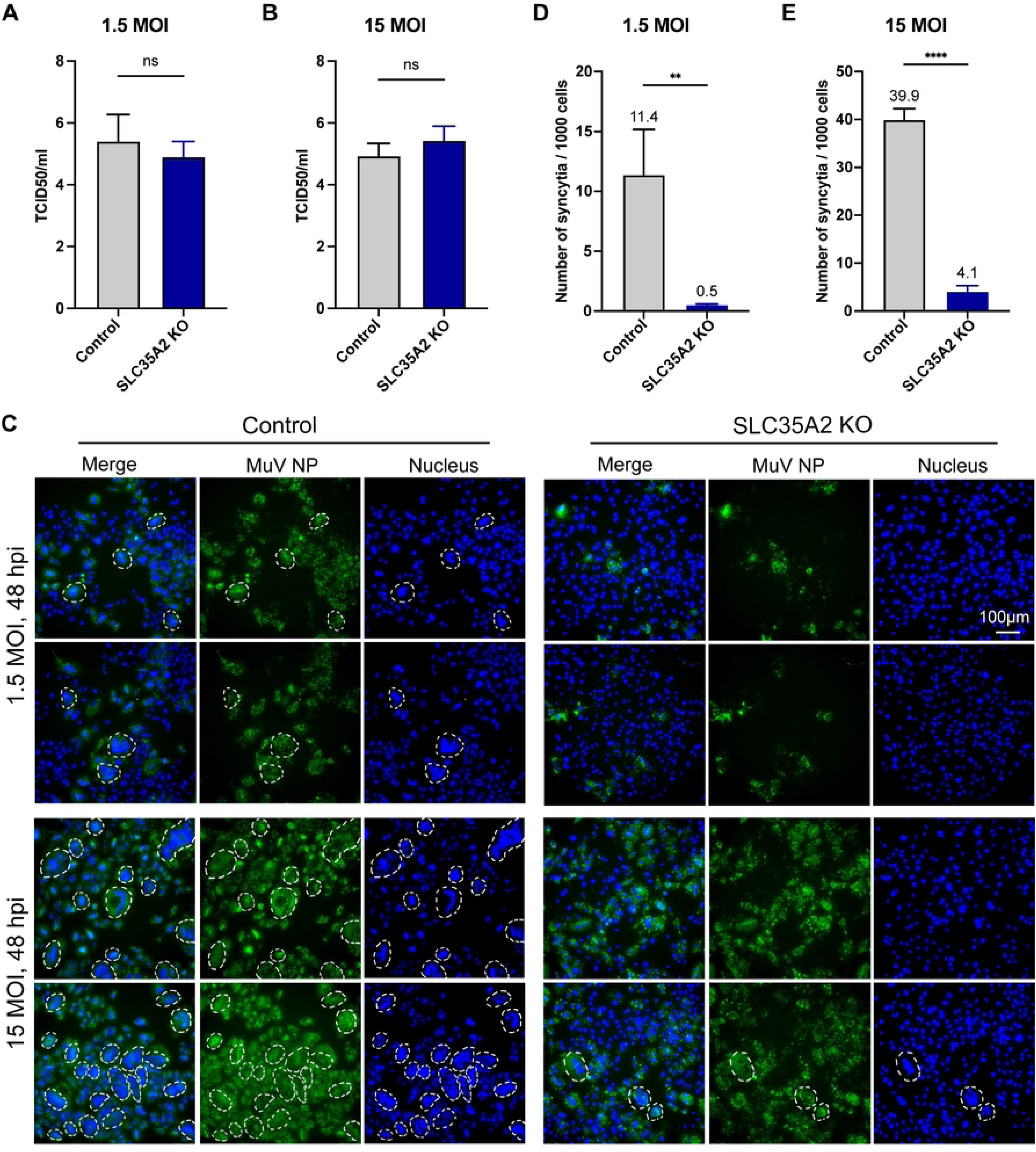
Measurement of MuV infectious particle production and MuV-induced syncytia. (A, B) MuV tissue culture infectious dose 50 (TCID_50_) from the supernatant of cells infected for 72 hours at an MOI of 1.5 MOI (A) or 15 (B). Data represent the mean of three independent experiments. Significance was calculated by unpaired t-test. ns: not significant. (C) Syncytia formation 48hpi with MuV at MOIs of 1.5 and 15. The white dashed lines mark the syncytia defined as MuV NP positive and with more than 3 nuclei. Data shown represent one of three independent experiments. (D and E) Number of syncytia among 1000 cells was counted at 48hpi at MOIs of 1.5 and 15. Data represent the mean of three independent experiments. Significance was calculated by unpaired t-test. **: p<0.01, ****: p<0.0001.

## Discussion

This manuscript presents the results of a CRISPR-Cas9 knockout screen identifying the transporters genes *SLC35A1* and *SLC35A2* as important for paramyxovirus infection and the characterization of their roles during paramyxovirus infections. *SLC35A1* is essential for the attachment of SeV, NDV, and MuV to the cell due to its role in exposing the viral receptor sialic acid on the cell surface, similar to what has been described for influenza virus and porcine delta coronavirus [29, 31]. Interestingly, while the role of *SLC35A2* is assumed to be related to virus attachment as galactose is typically considered the sugar to which sialic acid is linked [4], we found that *SLC35A2* is not essential for virus attachment to the cell surface. Instead, *SLC35A2* is crucial for the fusion of SeV with the cells and for MuV induced cell-to-cell fusion and syncytia formation suggesting a specific role for this protein in fusion events during virus infection.

Virus-cell fusion is an important target for antiviral drug development. To date, research on the fusion process and anti-fusion strategies have mainly focused on the role the viral proteins F and HN play during viral infection [44, 45]. However, our understanding of the host factors involved in the processes of virus-cell and cell-to-cell fusion that occur during infection are limited to the described role of 25HC in interfering with NiV induced cell-to-cell fusion [46], and a the role of soluble N-ethylmaleimide-sensitive factor attachment protein receptor (SNARE) protein USE1 in the glycosylation and expression of MuV fusion protein [47]. To our knowledge, no host factors have been identified to affect paramyxovirus virus-to-cell fusion.

Specific N-glycans on the F protein of several paramyxoviruses are important for the fusion activity of the protein [48–50]. However, we show successful infection of *SLC35A2* KO and control cell lines with the same MuV virus stocks and production of similar levels of MuV infectious viral particles in *SLC35A2* KO and control cells indicating normal activity of the F protein during virus entry in these conditions. These data suggest that either different glycosylations or factors beyond F glycosylation are impacted by *SLC35A2* during MuV infection. In addition, *SLC35A2* has been identified as an HIV X4 strain-specific restriction factor in primary target CD4+ T cells [51] and lack of *SLC35A2* resulted in decreased influenza virus polymerase activity using a viral replicon system [52]. However, our SeV and MuV genome replication data suggests *SLC35A2* has no effect on viral replication and transcription during infection with these viruses.

Sialic acids are well studied in influenza virus infections since sialic acids on cell surface glycoproteins and glycolipids serve as receptors for the influenza virus. Loss of *SLC35A1* causes reduced or abolished levels of sialylation on the cell surface, resulting in a severe impairment of influenza virus docking and entry [40, 52]. The role of *SLC35A2*, in contrast, is less clear. One study showed that absence of *SLC35A2* abolishes influenza H1N1 replication [53] but another study showed no effect on influenza H7N9 virus binding and internalization [52]. As we have confirmed that *SLC35A2* affects paramyxovirus virus-cell and cell-to-cell fusion, there is a possibility that this protein also affects fusion processes in infections with influenza and other viruses.

Interestingly, SeV could bind to *SLC35A2* KO cells and both NDV and MuV infected *SLC35A2* KO cells better than *SLC35A1* KO cells. These observations coupled with data from lectin staining showing that while *SLC35A2* KO deleted all surface galactose it only decreased cell surface sialic acid expression levels, suggest that while sialic acid is essential for the binding of these viruses to cells, galactose is not, indicating these viruses may utilize alternative sialic acid receptors beyond the classic sialic acid linked to galactose. Articles describing sialic acids as a receptor for viruses focus on sialic acid linked to galactose [4, 12, 13, 40, 54–59] but alternative linkages, such as α2,6-linkage to N-acetylgalactosamine are reported [4, 60]. Based on the data reported here, we hypothesize that either α2,6-linkage to N-acetylgalactosamine or other sialic acid linked molecules contribute to virus entry into A549 cells. Although further research is needed, our work provides new insights to the field.

Syncytia formation occurs late in the MuV life cycle when the two envelope proteins expressed on infected cells mediate fusion with neighboring cells [61]. Viral components can be transmitted between the fused infected cells through syncytia. Here, we observed that the MuV F mRNA level in *SLC35A2* KO cells was significantly lower at low MOI. Additionally, the amount of infectious viral particles produced in both cell lines was the same, suggesting that the lower level of viral mRNA is likely related to the reduced syncytia formation in *SLC35A2* KO cells. This finding suggests that *SLC35A2* promotes MuV cell-to-cell transmission through MuV induced syncytia formation.

In this work, we used the reporter virus rSeVC^dseGFP^ for screening. Compared to GFP, dseGFP, with a half-life of only two hours [36, 37], has significant advantages in CRISPR screening as it better reflects real-time viral replication activity. The identification of *SLC35A1* and *SLC35A2* in the dsGFP negative cell population indicates that our screening method is very effective. Here, we only show host factors identified from dseGFP negative cell population. Future work will focus on dseGFP low and high cell populations, where we aim to identify polymerase-related host factors, including those involved in non-standard viral genome generation.

In conclusion, our CRISPR-Cas9 KO screen identified *SLC35A1* and *SLC35A2* as critical host factors for paramyxovirus infection. *SLC35A1* is essential for viral binding, while *SLC35A2* is crucial for SeV-cell fusion and implicated in MuV-induced cell-to-cell fusion. Our findings show that even without *SLC35A2*, SeV can bind, NDV can enter and express genes, and MuV can complete its life cycle, indicating that galactose is not essential for viral attachment. This suggests the existence of alternative sialic acid receptors. Also, the reduced syncytia formation in *SLC35A2* knockout cells at low MOI suggests that *SLC35A2* deletion may inhibit MuV cell-to-cell transmission. These insights highlight the potential of targeting *SLC35A2* for therapeutic interventions against paramyxovirus infections.

## Materials & methods

### Cell lines

A549 cells (ATCC, #CCL-185), BSR-T7/5 cells (kindly provided by Dr. Conzelmann) [62], Lenti-293T cells (TaKaRa, # 632180), and LLC-MK2 cells (ATCC, #CCL-7) were cultured in tissue culture medium (Dubelcco’s modified Eagle’s medium (DMEM) (Invitrogen, #11965092) supplemented with 10% fetal bovine serum (FBS) (Sigma, #F0926), gentamicin 50 ng/ml (ThermoFisher, #15750060), L-glutamine 2 mM (Invitrogen, #G7513) and sodium pyruvate 1 mM (Invitrogen, #25-000-C1) at 5% CO_2_ 37℃. Cells were treated with mycoplasma removal agent (MP Biomedical, #3050044) and tested monthly for mycoplasma contamination using the MycoAlert Plus mycoplasma testing kit (Lonza, #LT07-318).

### Genetically modified cell lines

Lentiviruses for transduction were generated by co-transfecting a plasmid expressing the gene of interest or sgRNA together with the lentivirus packaging plasmids psPAX2 (Addgene, #12260) and pMD2.G (Addgene, #12259) into Lenti-293T cells using a TransIT-Lenti Transfection Reagent (Mirus Bio, # MIR 6604). Supernatants were collected 48 hours post transfection and lentivirus were detected by a Lenti-X™ GoStix™ Plusx kit (TaKaRa, #631280). A549 cells were then transduced with 500ul supernatants containing the lentivirus and 8 ug/ml polybrene (1200rpm, 30℃, 2 hours). The cells were transferred to 6 well plates the second day followed by antibiotic selection (Blasticidin 10 ug/ml for 1 week, Puromycin 0.5 ug/ml for 1 week, Hygromycin 400ug/ml for 2 weeks). Surviving cells were single cell cloned and confirmed by western blot or lectin staining.

A549-Cas9 stable line were generated by transducing lentiCas9-Blast plasmid (Addgene, #52962) [63] into A549 wt cells followed by Blasticidin selection and single cell cloning. A549-*SLC35A1*, A549-*SLC35A2* KO, and A549 control cell lines were made by transduction of sgRNA plasmids (Puromycin) to A549-Ca9 stable cell line. sgRNA plasmids with Puromycin resistance for *SLC35A1* (sgRNA: CCATAGCTTTAAGATACACA), *SLC35A2* (sgRNA: TGCGGGCGTAGCGGATGCTG) or a scrambled sgRNA control (CACTCACATCGCTACATCA) were ordered from Applied Biological Materials Inc. The *SLC35A1* and *SLC35A2* double KO cell line was made by transduction of a *SLC35A2* sgRNA plasmid (Hygromycin) to A549-*SLC35A1* KO cells.

### Viruses and virus infection

SeV Cantell expressing the reporter miR670 (rSeVC^miRF670^ [64]), SeV expressing HA-tagged M (rSeV-HA-M [43]) and the NDV reporter virus rNDV^eGFP^ were grown in 10-day-old, embryonated chicken eggs (Charles River) for 40 hours as previously described [65]. The VSV reporter virus (rVSV-eGFP) [66, 67] was obtained from Dr. Sean Whelan (Washington University in St.Louis) and grown in BSR-T7/5 cells. MuV (ATCC, #VR-1379) was grown in LLC-MK2 cells.

Infections with SeV, NDV, or MuV were performed by washing the cells once with PBS and incubating with virus diluted in infection media (DMEM, 35% bovine serum albumin (Sigma, #A7979), penicillin-streptomycin (Gibco, # 15140-122), and 5% NaHCO_3_ (Gibco, #25080094) at 37℃ for 1 hour, shaking every 15 minutes. Cells were then washed twice with PBS and supplemented with additional infection media. The infected cells were incubated at 37℃ until harvest.

### Recombinant reporter viruses rescue

The reporter viruses rSeVC^eGFP^ and rSeVC^dseGFP^ were rescued using the SeV Cantell strain reverse genetic system as described before [64]. First, two full-length plasmids pSL1180-rSeV-C^eGFP^ and pSL1180-rSeV-C^dseGFP^ were made by replacing the miRF670 gene of pSL1180-rSeV-C^miRF670^ with an eGFP or a destabilized eGFP (dseGFP) gene [36]. Additional nucleotides were inserted downstream of the dsGFP gene to ensure that the entire genome followed the “rule of six”. The viruses were rescued by co-transfecting full-length plasmids and the three helper plasmids to BSR-T7/5 cells using Lipofectamine LTX with Plus Reagent (Invitrogen, #15338100). The expression of GFP or dsGFP was monitored daily using fluorescence microscopy. At 4 days post-transfection, the cell cultures were harvested, and the supernatants were used to infect 10-day-old specific-pathogen-free embryonated chicken eggs via the allantoic cavity after repeated freeze-thaw cycles. After incubation for 40 hours at 37℃, 40-70% humidity, the allantoic fluids were harvested and the TCID_50_ was measured using LLC-MK2 cells.

The reporter virus rSeVC^GFP-ΔFHN+GFtail^ virus was generated and rescued by replacing SeV F and HN gene with a VSV G gene while retaining the SeV F protein tail. In brief, a VSV-GFtail plasmid was made by replacing G tail (G protein 490-511aa) with SeV F tail (F protein 524-565aa) using plasmid pMD2.G (Addgene, #12259), then the pSL1180-rSeVC^GFP-ΔFHN+GFtail^ full-length plasmid was made by cloning GFtail and deleting SeV F and HN gene from pSL1180-rSeVC^eGFP^ through PCR and In-Fusion cloning (TaKaRA Bio, #638948). Virus was rescued as described above, and after five serial passages on A549-*SLC35A1* KO cells, virus titer increased from 10^2^ to above 10^7^ TCID_50_/mL. Sanger sequencing confirmed the rescued virus sequence but revealed a D99G mutation within the M protein.

### Cas9 expression assay

Protein was extracted from A549-Cas9 stable cell line pool and cloned using 1% NP40 lysis buffer as described previously [64]. After a 20-minute incubation on ice and high-speed centrifugation for 20 minutes at 4℃, supernatant was collected, and protein concentration was quantified using the Pierce BCA Protein Assay Kit following the user’s guidelines (Thermo Fisher, # 23225). Next, 30 ug protein was denatured for 5 minutes at 95℃, loaded in a 4% to 12% Bis Tris gel (Bio-Rad, #3450124), and transferred to a PVDF membrane (Millipore Sigma, #IPVH00010). After blocking with 5% milk, membranes were incubated overnight with anti-CRISPR-Cas9 (Abcam, #ab191468) or anti-GAPDH (Sigma, #G8795) antibodies diluted in 5% BSA containing TBS with 0.1% Tween20. Membranes were incubated with anti-mouse secondary antibody conjugated with HRP for 1 hour in 5% BSA in TBST. Membranes were developed using Lumi-light western blotting substrate (Roche, #12015200001) and HRP was detected by a ChemiDoc (Bio-Rad).

### Cas9 activity assay

A549-Cas9 single cell clones were transduced with pXPR_011 lentivirus at an MOI of ∼1.0 in 12 well plates. pXPR_011 plasmid was a gift from John Doench & David Root [34] (Addgene, #59702). Transduced cells were transferred to 6 well plates on day 3 post transduction and treated with puromycin. On day 9 post transduction, surviving cells from each single cell clone were collected and eGFP signal was detected by spectral flow cytometry. Active Cas9-expressing lines resulted in a reduction of eGFP when transduced with pXPR_011 as this vector delivers both eGFP and a sgRNA targeting eGFP. Because eGFP is linked to puromycin gene with a 2A site, abrogation of eGFP will have no impact on puromycin resistance. The lower eGFP percentage of a single cell clone indicates higher Cas9 activity.

### A549-Brunello CRISPR KO library

A549-Cas9 stable cells were transduced at a low MOI (∼0.3) with Human CRISPR Knockout Pooled Library Brunello (Addgene, #73178) [35]. Transduction conditions and antibiotic concentration were optimized for the A549-Cas9 stable cell line (cell seeding density: 8 x 10^4^ per well (6-well plate); Puromycin concentraction: 0.5ug/ml. Lentivirus library was tittered to achieve a 30-50% infection rate and transductions were performed with 1.35×10^8^ cells to achieve a representation of at least 500 cells per sgRNA per replicate. Puromycin was added 2 days post transduction and was maintained for 5 - 7 days. The library cells containing sgRNA were used for CRISPR screening. Throughout the screen, the cells numbers were maintained at over 4×10^7^ cells to ensure coverage of at least 500 cells per sgRNA.

### CRISPR screening

A549-Brunello library was infected by SeV reporter virus rSeVC^dseGFP^ nsVG negative stock at an MOI of 10 or rSeVC^dseGFP^ nsVG positive stock at a MOI of 3 and cells were harvested followed by cell sorting to isolate the eGFP negative cell population. After screening, the sorted cells and two aliquots of the original CRISPR library (Mock) were pelleted and frozen at -80℃. Genomic DNA (gDNA) was isolated using the QIAamp DNA Blood Midi (Qiagen, #51183) or QIAamp DNA Bloop Mini (Qiagen, #51104) kit according to the manufacturer’s instructions. The concentration of each gDNA was measured on a Qubit Fluorimeter with the Qubit dsDNA Quantification Assay Kit (Invitrogen, #Q32851). CRISPR sgRNA barcodes were amplified by PCR of up to 10 μg of each gDNA template with a P5 stagger forward primer and P7 barcoded reverse primer [68]. In addition to gDNA, each 100 μL reaction contained 4 μL Titanium Taq DNA Polymerase (Takara, #639242), 10 μL Titanium PCR buffer, dNTPs at a final concentration of 100 μM, 5 μL DMSO (Sigma, #D9170-5VL), and P5 forward and P7 reverse primers each at a final concentration of 1 μM. PCRs were run for 5 minutes at 95℃, followed by 29 cycles of 95℃ for 30 s, 59℃ for 30 s, and 72℃ for 20 s, and a final extension step at 72℃ for 10 minutes. PCR amplicon concentrations were quantified on a Qubit Fluorimeter with the Qubit dsDNA Quantification Assay Kit before pooling and purifying with Agencourt AMPure XP SPRI beads (Beckman Coulter, #A63880) according to the manufacturer’s instructions. The final pool was submitted to the DNA Sequencing Innovation Lab (Washington University School of Medicine) for sequencing on the Illumina NextSeq-Mid platform with a 15% spike-in of PhiX DNA, yielding a total of 109,525,309 reads (Table 1). After demultiplexing according to the barcode sequences, the genes enrichment between cell populations was analyzed by MAGeCK (Version 0.5.9). The output tables were loaded and visualized with Prism 10.

**Table 1.** CRISPR screening read counts per sample.

| Sample | Read count |
| --- | --- |
| nsVG negative stock | 21,778,516 |
| nsVG negative stock + Ruxolitinib | 22,690,825 |
| nsVG positive stock | 22,583,165 |
| Mock | 60,290,626 |

### Lectin staining

Cells were seeded at a confluency of 1 × 10^5^ cells/well in a 12-well plate a day prior to staining. The next day, the cells were washed twice with PBS and fixed using 2% PFA at RT for 15 minutes. Following fixation, the cells were blocked with 3% BSA (in PBS) at RT for 1 hour. Lectin SNA-CY3 (VectorLabs, #CL-1303-1) or ECL-Fluorescein (VectorLabs, #FL-1141-5) was diluted in PBS at a 1:500 dilution and incubated with the cells on ice for 1 hour. The nuclei were then stained with a 1:10,000 dilution of Hoechst 33342 (Invitrogen, # H3570) at RT for 10 minutes.

### RNA extraction and RT-qPCR

Total RNA of infected cells and control samples were extracted using Kingfisher and a MagMAX™ mirVana™ Total RNA Isolation Kit (Thermo Fisher, #A27828) following manufacturer’s guidelines. 300-500ng of total RNA was used for cDNA synthesis with high-capacity RNA to cDNA kit (Thermo Fisher, #18080051). qPCR was performed using SYBR green (Thermo Fisher, # S7564) and 5 μM of reverse and forward primers (Table 2) for SeV NP, NDV NP, MuV F, and VSV NP genes on an Applied Biosystems QuantStudio 5 machine. Primers used for qPCR are listed in Table 2. Relative copy numbers were normalized to human *GAPDH* and human *β -actin* expression as described previously [69].

**Table 2.**
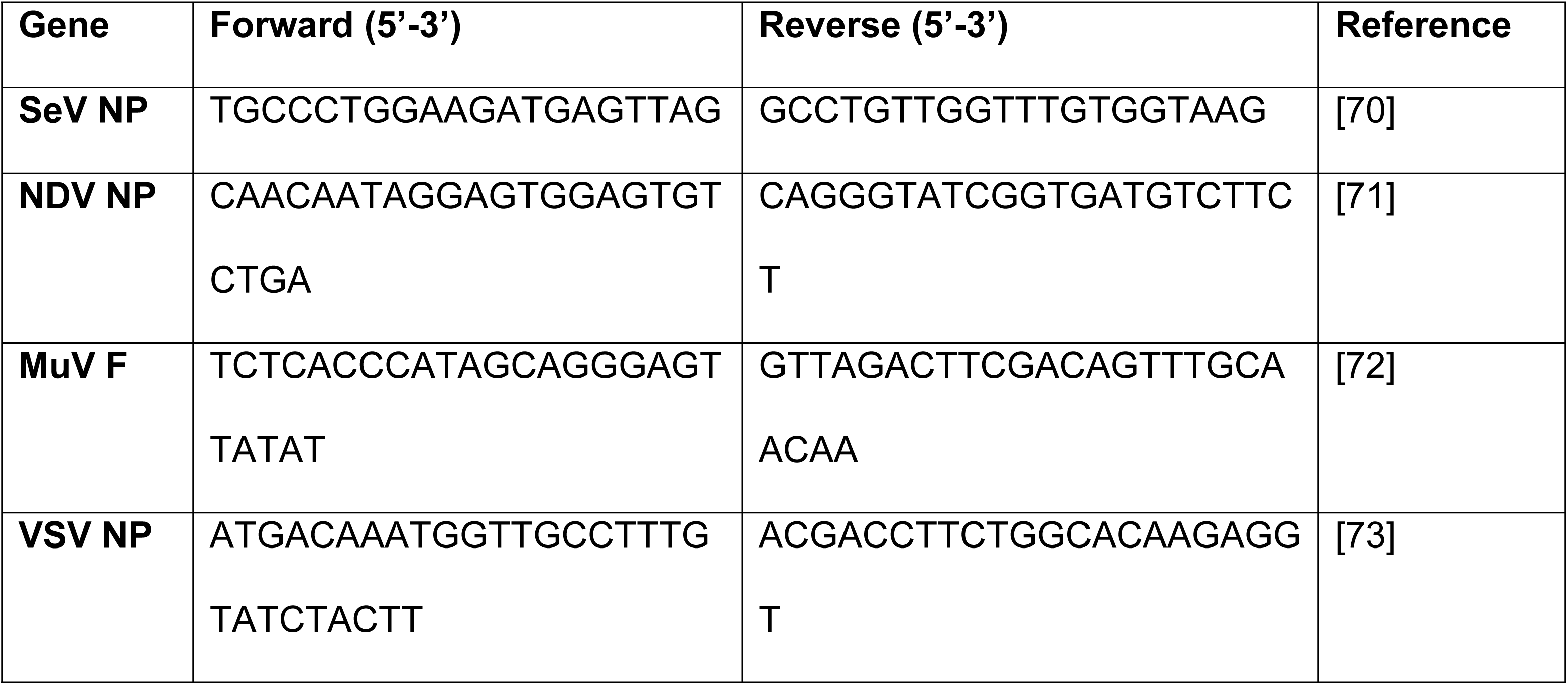
Primers for qPCR.

### *SLC35A1* and *SLC35A2* complementation

To complement *SLC35A1* or *SlC35A1* to KO cells, KO cell lines were transduced with the *SLC35A1*-GFP or *SLC35A2*-GFP cDNA expression constructs. In brief, *SLC35A1*-GFP and *SLC35A2*-GFP were obtained from plasmids pEGFP.N3-*SLC35A1*-GFP and pEGFP.N3-*SLC35A2*-GFP (Addgene 186281 and Addgene 186284)[74]. We performed codon optimization for the sgRNA binding sites of the target genes and switched to pLenti-Hygro plasmid backbone pLenti CMV Hygro DEST (Addgene, 17454) for lentivirus packaging. Then, *SLC35A1*-GFP and *SLC35A2*-GFP were separately introduced into their KO cell lines using the lentiviral transduction system. Following hygromycin selection, cell lines expressing the complemented genes were obtained, namely A549-*SLC35A1* KO+A1-GFP and A549-*SLC35A2* KO+A2-GFP. Subsequently, we infected these two cell lines with SeV reporter virus rSeVC^miRF670^ at a MOI of 3 and observed viral replication using fluorescence microscopy.

### NDV induced cell death quantification

To quantitatively analyze NDV-induced cell death, cells infected with rNDV-eGFP were collected at 24 and 48 hpi and analyzed by Cytek flow cytometry. In brief, to collect all the cells including cell debris and dead floating cells, debris and dead floating cells in supernatant were collected by spin down. The attached cells were collected after trypsinization. Then cells from each condition were merged and stained with eBioscience™ Fixable Viability Dye eFluor™ 506 (ThermoFosher, #65-0866-14) in 1:400 dilution on ice for 10 minutes. Cells are fixed with 2% PFA for 10 minutes at room temperature followed by flow analysis.

### MuV immunofluorescence

The infected cells were fixed using 2% PFA at RT for 15 minutes at specific time points post-infection followed by permeabilizing with 0.2% Triton X-100 (Sigma-Aldrich, # X100) for 10 minutes. Anti-MuV NP antibody (Thermo Fisher, #6008) was diluted in PBS at a 1:500 dilution and incubated at RT for 1 hour. Secondary antibodiy was diluted in PBS at a 1:500 dilution and incubated at RT for 30 minutes. The nuclei were stained with a 1:100,000 dilution of Hoechst 33342 (Invitrogen, # H-3570) along with the secondary antibody.

### Sendai virus binding experiment

To determine the binding capability of the Sendai virus to KO cells, we performed a virus-cell binding assay. Briefly, KO cells or control cells were incubated with the virus at a MOI of 30 at 4℃ for one hour. Following this, the cells were fixed with 4% paraformaldehyde (PFA) (Fisher Scientific, #50-980-495) for 10 minutes, blocked with 3% BSA for 30 minutes, and then stained with a HN Monoclonal Antibody-Alexa Fluor™ 647 (Thermo, # 51-6494-82) for 30 minutes. Finally, the results were analyzed using Cytek flow cytometry. During the procedure, cells were washed in PBS supplemented with 2% BSA and 2 mM EDTA (Corning, # 46-034-CI) 3 times between each step.

### Sendai virus Fusion experiment

To detect whether SeV underwent membrane fusion with the cell membrane, we referred to the Ebola membrane fusion experiment [42]. In brief, control cells or KO cells were incubated with 400 MOI rSeV-M-HA at 37℃ for 3 hours, then treated with 0.5mg/ml proteinase K (NEB, #P8107S) at 37℃ for 70 minutes to remove virus bound to the cell surface but not fused with the cells. Next, the samples were fixed and permeabilized with eBioscience™ Foxp3 / Transcription Factor Staining Buffer Set (Invitrogen, #00552300). The intracellular M-HA protein was then detected by a HA-PE mAb (Biolegend, #901518). SeV HN was detected using a HN Monoclonal Antibody-Alexa Fluor™ 647 (Thermo, #51-6494-82) as a control for cell surface proteins. Finally, the results were analyzed by Cytek flow cytometry. The detection of M protein in the proteinase K-treated group indicates that the virus particles have fused with the cells.

### MuV infectious particles measurement by TCID_50_

96-well plates were prepared the day before the experiment by seeding 20,000 A549wt cells per well the day before titration. The collected samples were serially diluted 10-fold in infection media, ranging from 1:10 to 1:10^8^. The plated cells were washed once with PBS, and 100 µL of each dilution was added to the cells, with each sample tested in triplicate. After incubating in a 37℃ incubator for 4 days, the cells were stained using the immunofluorescence method described above. The TCID_50_/ml was calculated based on the fluorescence results.

### MuV-induced syncytia quantification

After immunofluorescence staining, 3 images of each sample were captured using an inverted fluorescence microscope at both 20x and 5x magnifications. The 5x images were used for quantifying the MuV-induced syncytia. Fiji software was used to count the total number of nuclei in each 20x image, and the number of syncytia was manually marked and counted. Syncytia were defined as MuV-NP positive giant cells containing more than three nuclei. The number of syncytia per 1000 cells was then calculated. The results included three biological replicates.

### Statistics

Statistics were calculated using GraphPad Prism Version 10 (GraphPad Software, San Diego, CA).

## Acknowledgments

We thank Drs. Karl-Klaus Conzelmann for providing the BSR-T7/5 cells, Sean Whelan for the rVSV^eGFP^ virus, Susan Weiss for the rNDV^eGFP^ virus, and Benhur Lee for the rSeV-M-HA virus. We also acknowledge the funding for this project: C.L.B. was supported by NIH R01AI137062 and the BJC Investigator program at WUSTL. M.T.B. was supported by The G. Harold and Leila Y. Mathers Foundation and the Burroughs Wellcome Fund Pathogenesis of Infectious Disease Program. D.E.C. was supported by NIH T32 DK077653-29 and Crohn’s & Colitis Foundation Research Fellowship Award #935619.

## Supporting information

**S1 Fig. A549-Cas9 Stable Cell Line.** (A) The Cas9 expression of 6 A549-Cas9 single cell clones, A549wt cells, and A549-Cas9 pool was detected by western bolt. GAPDH expression was detected as a total cell protein control. (B) Diagram of Cas9 activity assay: with Cas9, GFP, and sgRNA targeting GFP in the same cells, higher Cas9 activity leads to lower GFP intensity, and lower Cas9 activity leads to higher GFP intensity. (C) Percentage of GFP-positive cells detected by flow cytometry after transduction of the pXPR_011 plasmid into A549wt cells and A549-Cas9 single cell clones. A549wt without transduction was used as a negative control. The red dashed line indicates 30% of GFP positive cells; it is generally accepted in the field that Cas9 cells with less than 30% GFP-positive cells can be used for CRISPR screening.

**S2 Fig. SeV Reporter Viruses rSeV^eGFP^ and rSeV^dseGFP^.** (A) Schematic representation of the SeV reporter viruses. eGFP or dseGFFP gene was inserted into the SeV genome between NP and P gene. dseGFP is made by fusion of a PEST degradation sequence to the C terminal of eGFP. (B) Fluorescence images of A549wt cells infected with rSeV^eGFP^ and rSeV^dseGFP^. A549wt cells were infected with an MOI of 3 of rSeV^eGFP^ or rSeV^dseGFP^ and images were analyzed at 24hpi. The nucleus was stained with Hoechst 33342 (Blue), green fluorescence indicates viral infection eGFP or dseGFP expression and accumulation. The images display three different fields of view. Scale bar lengths are indicated.

**S3 Fig. *SLC35A1* or *SLC35A2* KO Cells Infected with SeV at High MOI.** Fluorescence images showing GFP expression in control, *SLC35A1* KO, and *SLC35A2* KO cells infected with rSeVC^eGFP^ or rSeVC^dseGFP^ at MOIs of 5, 20, or 100, 24hpi. Scale bar lengths are indicated.

**S4 Fig. Analysis of Cell Death Induced by NDV Infection.** (A) Flow cytometry analysis of cell debris and cells in control, *SLC35A1* KO and *SLC35A2* KO cells infected with rNDV^eGFP^ at an MOI of 1.5 and control mock cells at 24 and 48 hpi. The percentages of cell debris or cells are indicated within each plot. Data shown represent one of three independent experiments. (B) Quantification of cell debris percentages at 24 and 48 hpi. Data represent the mean of three independent experiments. Statistical significance is indicated as follows: ****p < 0.0001, ns = not significant. (C) Flow cytometry analysis of cell death of cells population from (A) in control, *SLC35A1* KO, *SLC35A2* KO, and mock-infected cells at 24 and 48 hpi, as indicated by Fixable Viability Dye eFluor™ 506 staining. Percentages of dead cells are indicated within each plot. Data shown represent one of three independent experiments. (D) Quantification of dead cells percentages at 24 and 48 hpi. Data represent the mean of three independent experiments. Statistical significance is indicated as follows: *p < 0.05, ns = not significant.

**S5 Fig. rSeVC^eGFPΔFHN+GFtail^ Rescue and Confirmation.** (A) Schematic representation of the rSeVC^eGFPΔFHN+GFtail^. rSeVC^eGFPΔFHN+GFtail^ was designed by replacing SeV F and HN gene with a GSV-G deleting its tail and fusing with SeV F tail on rSeVC^eGFP^. Red inverted triangle indicated a mutation on M protein. (B) Fluorescence microscopy images of *SLC35A1* KO cells infected with rSeVC^eGFPΔFHN+GFtail^ at MOIs of 1.5, 3, and 6. Images were analyzed at 24 hpi. The nucleus was stained with Hoechst 33342 (blue), and green fluorescence indicates viral infection, as shown by eGFP expression. rSeVC^eGFP^ and Mock-infected cells were used as controls. Scale bar lengths are indicated.

**S1 Table. CRISPR screening data (nsVG negative stock)**

**S2 Table. CRISPR screening data (nsVG negative stock + Ruxolitinib)**

**S3 Table. CRISPR screening data (nsVG positive stock)**

**S4: MuV syncytia number**

## References

1. Duprex WP, Dutch RE. Paramyxoviruses: Pathogenesis, Vaccines, Antivirals, and Prototypes for Pandemic Preparedness. J Infect Dis. 2023;228(Suppl 6):S390–S7. doi: 10.1093/infdis/jiad123. PubMed PMID: 37849400; PubMed Central PMCID: PMCPMC11009463.

2. Gazal S, Sharma N, Gazal S, Tikoo M, Shikha D, Badroo GA, et al. Nipah and Hendra Viruses: Deadly Zoonotic Paramyxoviruses with the Potential to Cause the Next Pandemic. Pathogens. 2022;11(12). Epub 20221125. doi: 10.3390/pathogens11121419. PubMed PMID: 36558753; PubMed Central PMCID: PMCPMC9784551.

3. Navaratnarajah CK, Generous AR, Yousaf I, Cattaneo R. Receptor-mediated cell entry of paramyxoviruses: Mechanisms, and consequences for tropism and pathogenesis. J Biol Chem. 2020;295(9):2771–86. Epub 20200116. doi: 10.1074/jbc.REV119.009961. PubMed PMID: 31949044; PubMed Central PMCID: PMCPMC7049954.

4. Stencel-Baerenwald JE, Reiss K, Reiter DM, Stehle T, Dermody TS. The sweet spot: defining virus-sialic acid interactions. Nat Rev Microbiol. 2014;12(11):739–49. Epub 20140929. doi: 10.1038/nrmicro3346. PubMed PMID: 25263223; PubMed Central PMCID: PMCPMC4791167.

5. Chang A, Dutch RE. Paramyxovirus fusion and entry: multiple paths to a common end. Viruses. 2012;4(4):613–36.

6. Cantin C, Holguera J, Ferreira L, Villar E, Munoz-Barroso I. Newcastle disease virus may enter cells by caveolae-mediated endocytosis. J Gen Virol. 2007;88(Pt 2):559–69. doi: 10.1099/vir.0.82150-0. PubMed PMID: 17251575.

7. Zhao R, Shi Q, Han Z, Fan Z, Ai H, Chen L, et al. Newcastle Disease Virus Entry into Chicken Macrophages via a pH-Dependent, Dynamin and Caveola-Mediated Endocytic Pathway That Requires Rab5. J Virol. 2021;95(13):e0228820. Epub 20210610. doi: 10.1128/JVI.02288-20. PubMed PMID: 33762417; PubMed Central PMCID: PMCPMC8437353.

8. Porotto M, Fornabaio M, Kellogg GE, Moscona A. A second receptor binding site on human parainfluenza virus type 3 hemagglutinin-neuraminidase contributes to activation of the fusion mechanism. J Virol. 2007;81(7):3216–28. Epub 20070117. doi: 10.1128/JVI.02617-06. PubMed PMID: 17229690; PubMed Central PMCID: PMCPMC1866072.

9. Russell CJ, Kantor KL, Jardetzky TS, Lamb RA. A dual-functional paramyxovirus F protein regulatory switch segment: activation and membrane fusion. J Cell Biol. 2003;163(2):363–74. doi: 10.1083/jcb.200305130. PubMed PMID: 14581458; PubMed Central PMCID: PMCPMC2173521.

10. Jardetzky TS, Lamb RA. Activation of paramyxovirus membrane fusion and virus entry. Curr Opin Virol. 2014;5:24–33. Epub 20140216. doi: 10.1016/j.coviro.2014.01.005. PubMed PMID: 24530984; PubMed Central PMCID: PMCPMC4028362.

11. Song Z. Roles of the nucleotide sugar transporters (SLC35 family) in health and disease. Mol Aspects Med. 2013;34(2-3):590–600. doi: 10.1016/j.mam.2012.12.004. PubMed PMID: 23506892.

12. Kubota M, Takeuchi K, Watanabe S, Ohno S, Matsuoka R, Kohda D, et al. Trisaccharide containing alpha2,3-linked sialic acid is a receptor for mumps virus. Proc Natl Acad Sci U S A. 2016;113(41):11579–84. Epub 20160926. doi: 10.1073/pnas.1608383113. PubMed PMID: 27671656; PubMed Central PMCID: PMCPMC5068328.

13. Kubota M, Matsuoka R, Suzuki T, Yonekura K, Yanagi Y, Hashiguchi T. Molecular Mechanism of the Flexible Glycan Receptor Recognition by Mumps Virus. J Virol. 2019;93(15). Epub 20190717. doi: 10.1128/JVI.00344-19. PubMed PMID: 31118251; PubMed Central PMCID: PMCPMC6639266.

14. Markwell MA, Svennerholm L, Paulson JC. Specific gangliosides function as host cell receptors for Sendai virus. Proc Natl Acad Sci U S A. 1981;78(9):5406–10. doi: 10.1073/pnas.78.9.5406. PubMed PMID: 6272300; PubMed Central PMCID: PMCPMC348754.

15. Markwell MA, Paulson JC. Sendai virus utilizes specific sialyloligosaccharides as host cell receptor determinants. Proc Natl Acad Sci U S A. 1980;77(10):5693–7. doi: 10.1073/pnas.77.10.5693. PubMed PMID: 6255459; PubMed Central PMCID: PMCPMC350135.

16. Sanchez-Felipe L, Villar E, Munoz-Barroso I. alpha2-3- and alpha2-6-N-linked sialic acids allow efficient interaction of Newcastle Disease Virus with target cells. Glycoconj J. 2012;29(7):539–49. Epub 20120807. doi: 10.1007/s10719-012-9431-0. PubMed PMID: 22869099; PubMed Central PMCID: PMCPMC7088266.

17. Kumar N, Sharma S, Kumar R, Tripathi BN, Barua S, Ly H, et al. Host-directed antiviral therapy. Clinical microbiology reviews. 2020;33(3):10.1128/cmr.00168-19.

18. Li G, De Clercq E. Overview of antiviral drug discovery and development: viral versus host targets. 2021.

19. Watanabe T, Kawaoka Y. Influenza virus–host interactomes as a basis for antiviral drug development. Current opinion in virology. 2015;14:71–8.

20. Watanabe T, Kawakami E, Shoemaker JE, Lopes TJ, Matsuoka Y, Tomita Y, et al. Influenza virus-host interactome screen as a platform for antiviral drug development. Cell host & microbe. 2014;16(6):795–805.

21. Li C-C, Wang X-J, Wang H-CR. Repurposing host-based therapeutics to control coronavirus and influenza virus. Drug discovery today. 2019;24(3):726–36.

22. Mei M, Tan X. Current strategies of antiviral drug discovery for COVID-19. Frontiers in Molecular Biosciences. 2021;8:671263.

23. Faisca P, Desmecht D. Sendai virus, the mouse parainfluenza type 1: a longstanding pathogen that remains up-to-date. Research in veterinary science. 2007;82(1):115–25.

24. Kolakofsky D, Le Mercier P, Nishio M, Blackledge M, Crepin T, Ruigrok RWH. Sendai Virus and a Unified Model of Mononegavirus RNA Synthesis. Viruses. 2021;13(12). Epub 20211209. doi: 10.3390/v13122466. PubMed PMID: 34960735; PubMed Central PMCID: PMCPMC8708023.

25. Walter MJ, Morton JD, Kajiwara N, Agapov E, Holtzman MJ. Viral induction of a chronic asthma phenotype and genetic segregation from the acute response. The Journal of clinical investigation. 2002;110(2):165–75.

26. Xu J, Sun Y, Li Y, Ruthel G, Weiss SR, Raj A, et al. Replication defective viral genomes exploit a cellular pro-survival mechanism to establish paramyxovirus persistence. Nature communications. 2017;8(1):799.

27. Castro ÍA, Yang Y, Gnazzo V, Kim D-H, Van Dyken SJ, Lopez CB. Murine Parainfluenza Virus Persists in Lung Innate Immune Cells Sustaining Chronic Lung Pathology. bioRxiv. 2023.

28. Mercado-López X, Cotter CR, Kim W-k, Sun Y, Muñoz L, Tapia K, et al. Highly immunostimulatory RNA derived from a Sendai virus defective viral genome. Vaccine. 2013;31(48):5713–21.

29. Wang X, Jin Q, Xiao W, Fang P, Lai L, Xiao S, et al. Genome-wide CRISPR/Cas9 screen reveals a role for SLC35A1 in the adsorption of porcine deltacoronavirus. Journal of Virology. 2022;96(24):e01626–22.

30. Wang J, Liu H, Yang Y, Tan Y, Sun L, Guo Z, et al. Genome-scale CRISPR screen identifies TRIM2 and SLC35A1 associated with porcine epidemic diarrhoea virus infection. International Journal of Biological Macromolecules. 2023;250:125962.

31. Han J, Perez JT, Chen C, Li Y, Benitez A, Kandasamy M, et al. Genome-wide CRISPR/Cas9 screen identifies host factors essential for influenza virus replication. Cell reports. 2018;23(2):596–607.

32. Yi C, Cai C, Cheng Z, Zhao Y, Yang X, Wu Y, et al. Genome-wide CRISPR-Cas9 screening identifies the CYTH2 host gene as a potential therapeutic target of influenza viral infection. Cell reports. 2022;38(13).

33. Carette JE, Guimaraes CP, Varadarajan M, Park AS, Wuethrich I, Godarova A, et al. Haploid genetic screens in human cells identify host factors used by pathogens. Science. 2009;326(5957):1231-5.

34. Doench JG, Hartenian E, Graham DB, Tothova Z, Hegde M, Smith I, et al. Rational design of highly active sgRNAs for CRISPR-Cas9–mediated gene inactivation. Nature biotechnology. 2014;32(12):1262–7.

35. Doench JG, Fusi N, Sullender M, Hegde M, Vaimberg EW, Donovan KF, et al. Optimized sgRNA design to maximize activity and minimize off-target effects of CRISPR-Cas9. Nat Biotechnol. 2016;34(2):184–91. Epub 20160118. doi: 10.1038/nbt.3437. PubMed PMID: 26780180; PubMed Central PMCID: PMCPMC4744125.

36. Li X, Zhao X, Fang Y, Jiang X, Duong T, Fan C, et al. Generation of destabilized green fluorescent protein as a transcription reporter. Journal of Biological Chemistry. 1998;273(52):34970–5.

37. He L, Binari R, Huang J, Falo-Sanjuan J, Perrimon N. In vivo study of gene expression with an enhanced dual-color fluorescent transcriptional timer. Elife. 2019;8:e46181.

38. López CB. Defective viral genomes: critical danger signals of viral infections. Journal of virology. 2014;88(16):8720–3.

39. Genoyer E, López CB. The impact of defective viruses on infection and immunity. Annual review of virology. 2019;6(1):547–66.

40. Han J, Perez JT, Chen C, Li Y, Benitez A, Kandasamy M, et al. Genome-wide CRISPR/Cas9 Screen Identifies Host Factors Essential for Influenza Virus Replication. Cell Rep. 2018;23(2):596–607. doi: 10.1016/j.celrep.2018.03.045. PubMed PMID: 29642015; PubMed Central PMCID: PMCPMC5939577.

41. Banning A, Zakrzewicz A, Chen X, Gray SJ, Tikkanen R. Knockout of the CMP–Sialic Acid Transporter SLC35A1 in Human Cell Lines Increases Transduction Efficiency of Adeno-Associated Virus 9: Implications for Gene Therapy Potency Assays. Cells-Basel. 2021;10(5):1259.

42. Carette JE, Raaben M, Wong AC, Herbert AS, Obernosterer G, Mulherkar N, et al. Ebola virus entry requires the cholesterol transporter Niemann-Pick C1. Nature. 2011;477(7364):340-3. Epub 20110824. doi: 10.1038/nature10348. PubMed PMID: 21866103; PubMed Central PMCID: PMCPMC3175325.

43. Genoyer E, Kulej K, Hung CT, Thibault PA, Azarm K, Takimoto T, et al. The viral polymerase complex mediates the interaction of viral ribonucleoprotein complexes with recycling endosomes during sendai virus assembly. MBio. 2020;11(4):10.1128/mbio.02028-20.

44. Palgen J-L, Jurgens EM, Moscona A, Porotto M, Palermo LM. Unity in diversity: shared mechanism of entry among paramyxoviruses. Progress in molecular biology and translational science. 2015;129:1–32.

45. Contreras EM, Monreal IA, Ruvalcaba M, Ortega V, Aguilar HC. Antivirals targeting paramyxovirus membrane fusion. Curr Opin Virol. 2021;51:34–47. Epub 20210927. doi: 10.1016/j.coviro.2021.09.003. PubMed PMID: 34592709; PubMed Central PMCID: PMCPMC8994020.

46. Liu SY, Aliyari R, Chikere K, Li G, Marsden MD, Smith JK, et al. Interferon-inducible cholesterol-25-hydroxylase broadly inhibits viral entry by production of 25-hydroxycholesterol. Immunity. 2013;38(1):92–105. Epub 20121227. doi: 10.1016/j.immuni.2012.11.005. PubMed PMID: 23273844; PubMed Central PMCID: PMCPMC3698975.

47. Liu Y, Katoh H, Sekizuka T, Bae C, Wakata A, Kato F, et al. SNARE protein USE1 is involved in the glycosylation and the expression of mumps virus fusion protein and important for viral propagation. PLoS Pathogens. 2022;18(12):e1010949.

48. Segawa H, Yamashita T, Kawakita M, Taira H. Functional analysis of the individual oligosaccharide chains of Sendai virus fusion protein. The journal of biochemistry. 2000;128(1):65–72.

49. McGinnes L, Sergel T, Reitter J, Morrison T. Carbohydrate modifications of the NDV fusion protein heptad repeat domains influence maturation and fusion activity. Virology. 2001;283(2):332–42.

50. Hu A, Cathomen T, Cattaneo R, Norrby E. Influence of N-linked oligosaccharide chains on the processing, cell surface expression and function of the measles virus fusion protein. Journal of general virology. 1995;76(3):705–10.

51. Itell HL, Humes D, Baumgarten NE, Overbaugh J. Host cell glycosylation selects for infection with CCR5-versus CXCR4-tropic HIV-1. bioRxiv. 2023.

52. Yi C, Cai C, Cheng Z, Zhao Y, Yang X, Wu Y, et al. Genome-wide CRISPR-Cas9 screening identifies the CYTH2 host gene as a potential therapeutic target of influenza viral infection. Cell Rep. 2022;38(13):110559. doi: 10.1016/j.celrep.2022.110559. PubMed PMID: 35354039.

53. Carette JE, Guimaraes CP, Varadarajan M, Park AS, Wuethrich I, Godarova A, et al. Haploid genetic screens in human cells identify host factors used by pathogens. Science. 2009;326(5957):1231-5. doi: 10.1126/science.1178955. PubMed PMID: 19965467.

54. Kuchipudi SV, Nelli RK, Gontu A, Satyakumar R, Surendran Nair M, Subbiah M. Sialic acid receptors: the key to solving the enigma of zoonotic virus spillover. Viruses. 2021;13(2):262.

55. Sieben C, Sezgin E, Eggeling C, Manley S. Influenza A viruses use multivalent sialic acid clusters for cell binding and receptor activation. PLoS pathogens. 2020;16(7):e1008656.

56. Zhao C, Pu J. Influence of host sialic acid receptors structure on the host specificity of influenza viruses. Viruses. 2022;14(10):2141.

57. Liu M, Huang LZ, Smits AA, Büll C, Narimatsu Y, van Kuppeveld FJ, et al. Human-type sialic acid receptors contribute to avian influenza A virus binding and entry by hetero-multivalent interactions. Nature communications. 2022;13(1):4054.

58. Sun X-L. The role of cell surface sialic acids for SARS-CoV-2 infection. Glycobiology. 2021;31(10):1245–53.

59. Nguyen L, McCord KA, Bui DT, Bouwman KM, Kitova EN, Elaish M, et al. Sialic acid-containing glycolipids mediate binding and viral entry of SARS-CoV-2. Nature Chemical Biology. 2022;18(1):81–90.

60. Monteiro RC, Moura IC, Launay P, Tsuge T, Haddad E, Benhamou M, et al. Pathogenic significance of IgA receptor interactions in IgA nephropathy. TRENDS in molecular Medicine. 2002;8(10):464–8.

61. Lamb RA, Parks GD. Paramyxoviridae. Fields Virology: Sixth Edition: Wolters Kluwer Health Adis (ESP); 2013.

62. Buchholz UJ, Finke S, Conzelmann K-K. Generation of bovine respiratory syncytial virus (BRSV) from cDNA: BRSV NS2 is not essential for virus replication in tissue culture, and the human RSV leader region acts as a functional BRSV genome promoter. Journal of virology. 1999;73(1):251–9.

63. Sanjana NE, Shalem O, Zhang F. Improved vectors and genome-wide libraries for CRISPR screening. Nature methods. 2014;11(8):783–4.

64. González Aparicio LJ, Yang Y, Hackbart M, López CB. Copy-back viral genomes induce a cellular stress response that interferes with viral protein expression without affecting antiviral immunity. PLoS biology. 2023;21(11):e3002381.

65. López CB, García-Sastre A, Williams BRG, Moran TM. Type I interferon induction pathway, but not released interferon, participates in the maturation of dendritic cells induced by negative-strand RNA viruses. The Journal of infectious diseases. 2003;187(7):1126–36.

66. Whelan S, Ball LA, Barr JN, Wertz G. Efficient recovery of infectious vesicular stomatitis virus entirely from cDNA clones. Proceedings of the National Academy of Sciences. 1995;92(18):8388–92.

67. Cherry S, Doukas T, Armknecht S, Whelan S, Wang H, Sarnow P, et al. Genome-wide RNAi screen reveals a specific sensitivity of IRES-containing RNA viruses to host translation inhibition. Genes & development. 2005;19(4):445–52.

68. Sanson KR, Hanna RE, Hegde M, Donovan KF, Strand C, Sullender ME, et al. Optimized libraries for CRISPR-Cas9 genetic screens with multiple modalities. Nat Commun. 2018;9(1):5416. Epub 20181221. doi: 10.1038/s41467-018-07901-8. PubMed PMID: 30575746; PubMed Central PMCID: PMCPMC6303322.

69. Garcia GL, Valenzuela A, Manzoni T, Vaughan AE, Lopez CB. Distinct Chronic Post-Viral Lung Diseases upon Infection with Influenza or Parainfluenza Viruses Differentially Impact Superinfection Outcome. Am J Pathol. 2020;190(3):543–53. Epub 20191219. doi: 10.1016/j.ajpath.2019.11.003. PubMed PMID: 31866346; PubMed Central PMCID: PMCPMC7073775.

70. Yount JS, Kraus TA, Horvath CM, Moran TM, López CB. A novel role for viral-defective interfering particles in enhancing dendritic cell maturation. The Journal of Immunology. 2006;177(7):4503–13.

71. Qiu X, Yu Y, Yu S, Zhan Y, Wei N, Song C, et al. Development of strand-specific real-time RT-PCR to distinguish viral RNAs during Newcastle disease virus infection. The Scientific World Journal. 2014;2014(1):934851.

72. Tipples G, Hiebert J. Detection of measles, mumps, and rubella viruses. Diagnostic Virology Protocols. 2011:183–93.

73. Wang B, Yang C, Tekes G, Mueller S, Paul A, Whelan SP, et al. Recoding of the vesicular stomatitis virus L gene by computer-aided design provides a live, attenuated vaccine candidate. MBio. 2015;6(2):10.1128/mbio.00237-15.

74. Li D, Mukhopadhyay S. Functional analyses of the UDP-galactose transporter SLC35A2 using the binding of bacterial Shiga toxins as a novel activity assay. Glycobiology. 2019;29(6):490–503. doi: 10.1093/glycob/cwz016. PubMed PMID: 30834435; PubMed Central PMCID: PMCPMC6521944.

